# Inference in wolves and dogs: The “cups task”, revisited

**DOI:** 10.1101/2024.09.03.610928

**Authors:** Dániel Rivas-Blanco, Sophia D. Krause, Sarah Marshall-Pescini, Friederike Range

## Abstract

Inferential reasoning —the process of arriving at a conclusion from a series of premises— has been studied in a multitude of animal species through the use of the “cups task” paradigm. In one of the versions of this set-up, two opaque cups —one baited, one empty— are shaken in front of the animal. As only the baited cup makes a noise when shaken, the animals can locate the reward by inferring that only a baited cup would make noise, that an empty cup would make no noise, or both. In a previous iteration of this paradigm in wolves (*Canis lupus*) and dogs (*Canis familiaris*), wolves seemed to outperform dogs. However, due to the lack of control conditions, it was not possible to assess each species’ inference capabilities, nor how they related to each other. The current study adds several conditions in which the baited cup, the empty cup, or no cups are shaken, in order to tackle this issue. Our results seem to indicate that wolves and dogs made their choices not based on inference but on the saliency and order of the stimuli presented, something that seems in line with the previous study. We discuss the potential causes behind the animals’ performance, as well as proposing alternative paradigms that may be more apt to measure inference abilities in wolves and dogs.

## Introduction

Animals are exposed to a variety of stimuli linked to different opportunities and challenges. In order to make decisions and act appropriately, they need to process the available information in their environment (Völter & Call, 2017). One of such process methods is inferential reasoning, the ability of deriving a conclusion from previously known information (Völter & Call, 2017). The effects of inferential reasoning are sometimes difficult to distinguish from associative learning (i.e., linking stimuli that covary). However, whereas associative learning involves associating events that are simultaneously occurring in space and time (and thus requires trial and error), reasoning combines perceived events with imagined ones or events that are not necessarily co-occurring (Maier & Schneirla, 1935, discussed in Völter & Call, 2017) by retrieving and recombining information (Zeithamova et al., 2012). Accordingly, we can distinguish inferences from associative learning based on the prior knowledge of the animal and whether or not trial-and-error learning occurs before arriving at the solution. Importantly, inferential reasoning is thought to be responsible for animals being able to act appropriately even when only partial information is available (Premack, 1995, discussed in Völter & Call, 2017; Völter & Call, 2017).

One of the types of inferential reasoning is diagnostic inference (i.e., reasoning from the consequent — the effect— to the antecedent —the cause—). This can, in turn, be based on affirming the consequent or denying the consequent. Choosing to approach a rustling bush by inferring there may be prey hidden inside would be “affirming the consequent”. Conversely, avoiding a still bush (because the animal infers that no prey is hidden in it) would be “denying the consequent”. The latter type of inferential reasoning has been shown much more rarely in non-human species (see Völter & Call, 2017 for a review).

One of the paradigms used to analyze inference strategies in animals is the “cups task”, in which subjects can choose between two opaque cups (one containing a reward, while the other remains empty) based on information about the contents of the cups. For example, when a baited cup is shaken, an auditory cue in the form of a rattling sound is produced, indicating the presence of the food reward. Choosing the baited cup by using this cue to infer the presence of its cause is an example of “affirming the consequent” (Völter & Call, 2017). In contrast, avoiding the cup that does not make a sound when shaken (and is thus supposedly empty), would be “denying the consequent” (Völter & Call, 2017).

Several species have been tested in some version of the “cups task”, African grey parrots (Schloegl et al., 2012), pigs (Nawroth & von Borell, 2015), and apes (Call, 2004; Call, 2006; Bräuer et al. 2006; Hill et al., 2011), with all of them demonstrating their ability to make diagnostic inferences (see Hill et al., 2011, for a review). Studies on domestic dogs using the cups paradigm, on the other hand, have shown that dogs seem to lack such causal understanding (Bräuer et al., 2006; Erdőhegyi et al., 2007). Lampe et al. (2017) compared dogs with wolves, their closest living relatives (Vila et al., 1997). Similar to the above-mentioned studies, dogs performed at chance level, whereas wolves were able to solve the task above chance (Lampe et al., 2017).

Indeed, the results of the study performed by Lampe and colleagues suggest that wolves may outperform dogs in causal tasks, which would hint to an effect of domestication on dogs’ causal understanding. However, Lampe and colleagues only tested one condition (the so-called “noise” condition), where both a baited and an empty cup were shaken. They also did not include any control conditions to eliminate other potential explanations —most notably, that wolves may have been following the most salient stimulus, regardless of whether they were able to infer that the reward was there. Furthermore, the cups were always shaken in the same order (with the baited cup being shaken first and the empty one being shaken second), which leaves open the possibility of wolves solving the task by staying by the side where a cup was first shaken (again, regardless of whether they were able to infer the presence of the food). Consequently, it is not possible to ascertain which strategy the wolves may have employed to solve this task (be it inference by affirming the consequent, denying the consequent, or both) or whether inferential reasoning was involved at all, since they may have chosen the ‘full cup’ simply because it represented the most salient stimulus (the one that produced a noise) or by choosing the very first object being moved (primacy effect).

Thus, the study at hand aimed to investigate the inferential reasoning strategies used by wolves and dogs by replicating the “noise” condition of Lampe et al. (2017) and adding further conditions. Aside from the “full information” condition (both cups, one baited and one empty, are shaken; as in Lampe et al., 2017), the following conditions were included: B) “affirming the consequent” (only baited cup is shaken), C) “denying the consequent” (only empty cup is shaken), and the control condition D) “no information” (no cup is shaken) (**Table 3**).

According to Frank’s (1980) information processing hypothesis, wolves should generally perform better than dogs on inferential reasoning tasks. This hypothesis states that selection pressures on independent problem-solving skills and causal understanding were relaxed in dogs during the domestication process as a result of humans providing direct (in the case of pet dogs) or indirect (e.g. refuse to feed on in case of free-ranging dogs) care (Frank, 1980; Frank & Frank, 1982; Frank & Frank, 1985; Frank & Frank, 1987, discussed in Wynne, 2021; see also feeding ecology hypothesis, Marshall-Pescini et al., 2017a). This is further compounded by wolves possibly relying on complex mental processes such as means-ends relationships, causal understanding, and strategy while hunting (Frank & Frank, 1985). Indeed, several studies have observed better problem-solving skills in wolves than in dogs, which might be explained by wolves’ better causal understanding, but could also come as a result of the higher persistence displayed by wolves in such tasks, leading to success by trial- and error learning (Frank & Frank, 1982; Frank & Frank, 1985; Frank, 2011; Hiestand, 2011; see Bensky et al., 2013 for a review; Udell, 2015; Marshall-Pescini et al., 2017b).

Taken together, we expect wolves to have a better understanding of causation than dogs, performing better than chance in both the “full information” (both cups shaken), and both partial information conditions (“affirming the consequent” and “denying the consequent”). However, if the wolves’ performance in Lampe and colleagues’ study was due to choosing the most salient stimulus (i.e., a saliency effect), we expect performances above chance level in the “full information” and “affirming the consequent” condition and below chance level in the “denying the consequent” condition. Moreover, if the subjects simply choose based on the last or first stimulus shaken (i.e., a recency effect or primacy effect), we instead expect them to choose the second or first shaken cup in the “full information” condition and the shaken cup in both partial information conditions above chance. Lastly, as no cues are given in the “no information” condition, we expect the subjects to perform at chance level if they do not use any alternative strategies (such as choosing based on olfactory cues). A detailed overview of the expected performance for each possible strategy the animals might have used within the different conditions is given in **Table 1**.

**Table 1:** Expected performance in each condition for the possible strategies the animals might use. . If “baited cup” or “empty cup” is stated in the table, this option is expected to be chosen above chance level. The bold outcome in the table shows the result in the respective condition which would be essential for differentiating between the possible strategies.

| Strategy | Full information | Partial information |  | No information |
| --- | --- | --- | --- | --- |
|  |  | Affirming the consequent | Denying the consequent |  |
| Causal understanding | Baited cup | Baited cup | **Baited cup<sup>1</sup>** | Random / side-bias |
| Most salient stimulus | Baited cup | Baited cup | **Empty cup** | Random / side-bias |
| Last stimulus | **Last cup shaken** | Baited cup | Empty cup | Random / side-bias |
| First stimulus | **First cup shaken** | Baited cup | Empty cup | Random / side-bias |
| Lack of understanding | Random / side-bias | Random / side-bias | Random / side-bias | Random / side-bias |

<sup>1</sup>: According to Völter & Call (2017), the ability to reason by denying the consequent is less widespread among animals than the ability to reason by affirming the consequent. Therefore, random/side bias results would also be compatible with this strategy.

## Material and Methods

### Subjects

In this study, we tested similarly raised wolves and dogs (hereafter “pack dogs”) at the Wolf Science Center (WSC) in Ernstbrunn, Austria. The wolves (*N*_wolf_ = 12; 8 males; mean age = 106.5 ± 11.2 months) and the pack dogs (*N*_pack dog_ = 5; 2 males, mean age = 98.4 ± 6.4 months) were hand-raised and had permanent contact to humans until the age of 5 months before being introduced or separated into packs. The wolves and pack dogs are living under similar conditions in outdoor enclosures measuring 2000 to 10.000 m^2^.

Based on previous studies (e.g., Müller et al., 2016), we had reason to believe that dogs raised with different experiences in regard to object manipulations and opportunities to learn physical properties of the environment should not differ in their inferential reasoning skills. Therefore, in order to increase our dog sample size, we recruited 7 additional pet dogs (*N*_pet dog_ = 7; 5 males; mean age = 99.3 ± 12.5 months). These pet dogs were familiar with the WSC test environment (so as to minimize potential differences in testing) with 4 of them being former pack dogs raised at the WSC. Nevertheless, in order to account for any potential differences in understanding of the contingencies of the task due to pet dogs having more experience with human-created objects, we compared the performance of both dog populations before merging their data for further analyses (and only did so after finding that, indeed, no significant differences arose between them, as explained in the Results section).

Three animals (2 wolves and 1 pet dog) were excluded due to motivational or health reasons (sight and hearing) during the training phase (see below) and thus not included in the statistical analyses. For the purpose of the analyses, therefore, the sample sample size for wolves was *N*_wolf_ = 10 and the one for pet dogs was *N*_pet dog_ = 6. Details about the subjects are provided in **Table 2**

**Table 2:** List of animals, indicating species, status, sex and age (at the time of the first training session). The horizontal division lines indicate the packs or households the animals live in together. Animals shown in grey and italics were excluded from the experiment (and thus, from any subsequent analyses) due to motivational or health problems. Please refer to **Table S1** for more details about the exact date of birth, exact date of the first training session, genetic relationships and breed information.

| Subject | Species | Status | Sex | Age (months) |
| --- | --- | --- | --- | --- |
| Chitto | Wolf | pack | m | 114 |
| *Tala* | *Wolf* | *pack* | *f* | *114* |
| Una | Wolf | pack | f | 115 |
| *Nanuk* | *Wolf* | *pack* | *m* | *150* |
| Taima | Wolf | pack | f | 66 |
| Tekoa | Wolf | pack | m | 66 |
| Geronimo | Wolf | pack | m | 150 |
| Yukon | Wolf | pack | f | 150 |
| Etu | Wolf | pack | m | 71 |
| Maikan | Wolf | pack | m | 71 |
| Kenai | Wolf | pack | m | 143 |
| Amarok | Wolf | pack | m | 119 |
| Hiari | Dog | pack | m | 92 |
| Imara | Dog | pack | f | 92 |
| Layla | Dog | pack | f | 124 |
| Panya | Dog | pack | f | 92 |
| Enzi | Dog | pack | m | 92 |
| Hakima | Dog | pet | m | 136 |
| *Kilio* | *Dog* | *pet* | *m* | *144* |
| Asali | Dog | pet | m | 135 |
| Freya | Dog | pet | f | 80 |
| Pepeo | Dog | pet | m | 93 |
| Zazu | Dog | pet | m | 93 |
| Coco | Dog | pet | f | 59 |

### Experimental conditions and set-up

#### Testing environment

The experiment was performed at the Wolf Science Center (WSC) in Ernstbrunn, Austria. The experimental sessions were mostly conducted in two large fenced outdoor enclosures, which are used for testing purposes. However, 5 wolves were tested in their home enclosure due to their unwillingness to leave that enclosure. In this case, the other pack members were shifted to another enclosure during the test.

#### Apparatus

The apparatus was a table (150 cm length, 50 cm width, 113-143 cm height —adjustable) positioned outside the enclosure behind the mesh-wire fence. The upper part of the table was surrounded by acrylic glass, through which the animals could observe the experimental procedure. Additionally, a curtain (96 x 57 cm) was attached behind the acrylic glass in the middle of the upper part of the table, concealing the experimenter and their actions to avoid providing unintended cues to the subject.

Two retractable targets, one on each side of the table, were used for the animals to indicate their choices in the experiment. Both targets were attached to a PVC pipe, enabling the experimenter to push them into the enclosure (within the animal’s reach) and back again. To ensure the targets were moved simultaneously, these two pipes were connected to each other by a third one (parallel to the long side of the table, perpendicular to the other two pipes) on the experimenter’s side of the table.

For the test trials, two containers –two plastic flower pots with a diameter of 14 cm– were used. The two cups were attached to wooden poles (45 cm in length) by two o-screws and two rubber bands, in such a way that the bands enveloped the cups while also being inserted into the circular cavity of the o-screws. This contraption allowed the experimenter to shake the cups from behind the curtain with a minimum amount of movement and without having their hands within the subject’s view (which could distract the subjects or allow them to receive unintentional cues from the experimenter).

To observe the subject’s position and choice despite the curtain, a GoPro Hero 4 camera installed at the top of the table provided the experimenter with a live video feed on a tablet during the test and filmed the session. In order to allow for double checking of the performance of the animals in each session at a later point, a second camera (Sony HDR) was placed outside the enclosure filming the session diagonally sideways from the animal’s point of view.

A detailed illustration of the testing table is shown in **Figure 1**.

**Figure 1:**
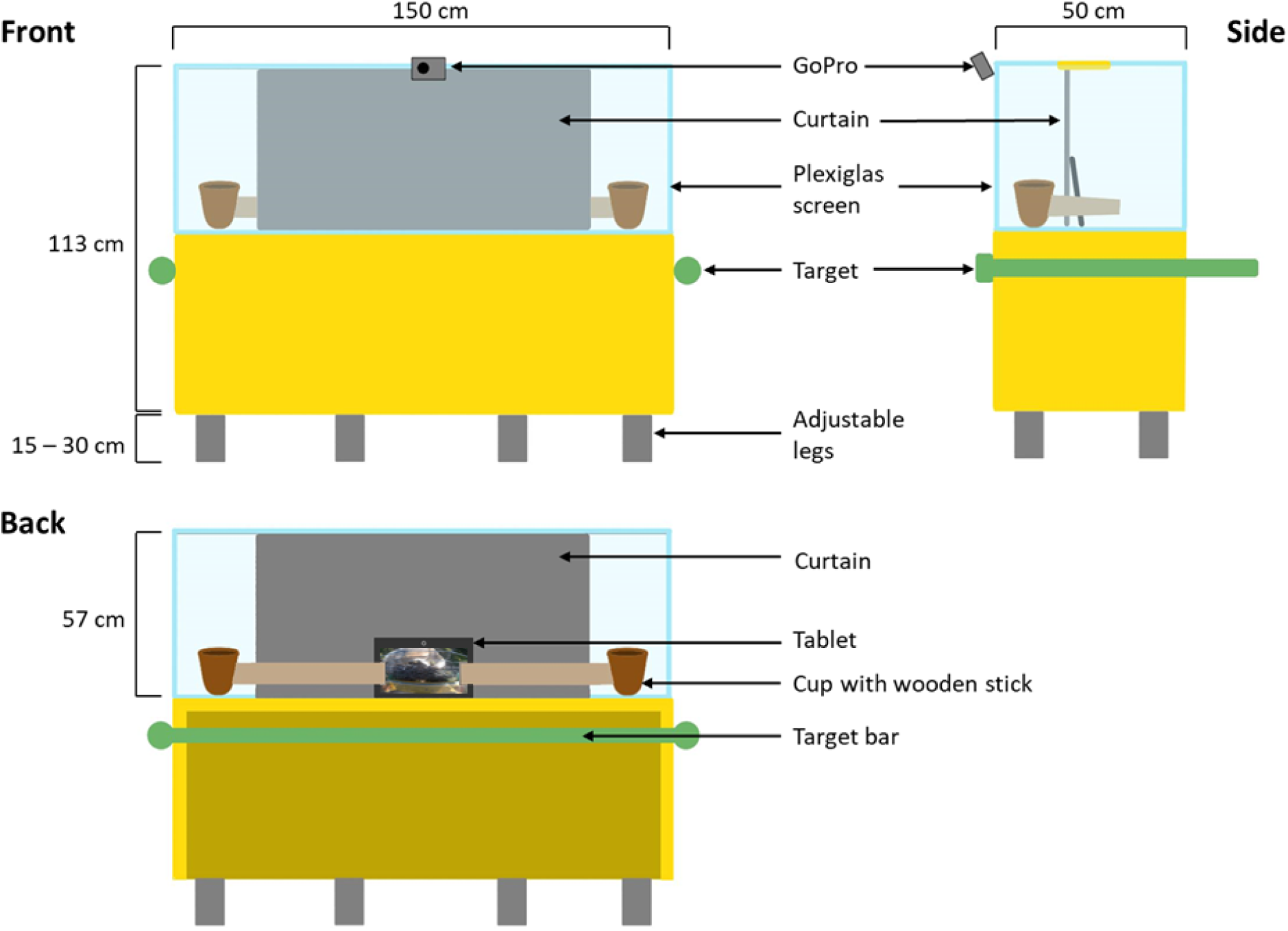
Schematic illustration of the testing apparatus. The front side represents the perspective of the animal. This side was positioned against the enclosure fence from the outside. The back side was the perspective of the experimenter who was sitting or standing behind the table. Yellow parts represent wooden parts of the table.

### Habituation

Three of the wolves that participated in this experiment (*Maikan, Taima, and Tekoa*) have shown to be particularly neophobic in other experiments and during daily activities. Thus, in order to reduce their fear towards the apparatus, a habituation phase preceded the experiment for these animals. In this phase, the animals were familiarized with the entire apparatus, especially the movable elements of it (moving targets, shaking cups and the noise they made). This was done to ensure that they could freely interact with the contingencies of the apparatus without fear. However, in order to make sure these animals did not have more experience on the relevant features of the task that may have given them an advantage over other subjects, we made sure never to allow them to interact directly with the cup nor see its contents. Furthermore, in order to prevent any potential associations from arising during this habituation phase, approaching both the empty and the baited cup was rewarded by the experimenter. The habituation phase lasted until the subject felt comfortable enough to interact with the table and participate in the test.

### Experimental procedure

#### General procedure

Prior to each training or testing day, the subjects were separated from their pack and shifted into the testing enclosure. This was followed by a short acclimatization period of approximately 5 minutes, in which the subjects had the opportunity to roam freely in the enclosure.

Each animal was tested in approximately 2-3 sessions per day on in 2-3 days a week. Each session had a time limit of 30 minutes. If the animal did not complete all trials within this time limit or showed no interest in the test (e.g. refused to be shifted into the test enclosure or did not approach the table), that session was cancelled and repeated on a different day. After 5 consecutive cancelled sessions (training or testing), a clear lack of motivation for the test, or the inability to participate in the test due to health reasons; the animal was excluded from the experiment. We excluded 2 of the wolves (*Tala* and *Nanuk*) and 1 pet dog (*Kilio*) in this manner.

#### Training phase

During this phase, the animals were trained to indicate their choice (“left” or “right”) by touching one of the targets on the side where the reward was placed. Each training session consisted of 12 trials. A training trial started by calling the animals’ attention by either saying their name out loud or knocking on the center of the table. Subsequently, a food reward (usually a small piece of beef or sausage, but chick pieces were used for some of the animals) was placed directly on one side of the table (outside of the area covered by the curtain), with the side on which the reward was placed (“left” or “right”) randomized and counterbalanced. This was followed by pushing the targets through the fence. If the subjects touched the target closest to the reward, the experimenter grabbed the reward and threw it over the fence and into the enclosure. Conversely, if the subject touched the target on the side opposite to the reward, the reward was instead taken back behind the curtain.

If the subjects chose the correct side in 10 out of 12 trials in two consecutive training sessions, they moved forward to the testing phase.

#### Testing phase

##### Pre-test

Prior to each test session, the subject was tested for motivation, attentiveness, and any side preferences in a *pre-test.* This *pre-test* consisted of 6 trials, identical to the training trials described above. The subjects had to choose the correct side at least 5 out of those 6 trials in order to proceed with the experimental trials. If this criterion was not reached, the animal got five more minutes of acclimatization and the *pre-test* was repeated. However, if the subject did not reach the criterion in two *pre-tests* on the same day, the test was cancelled for that day and rescheduled. After three consecutive failed *pre-tests*, the animal went back to the training phase (this happened only once with one of the pack dogs; *Layla*).

##### Experimental trials

Once the subject successfully completed the *pre-test*, they proceeded with the experimental trials. Each animal was tested in 8 test sessions, each consisting of 8 trials. Before a trial started, the experimenter prepared the cups behind the curtain by baiting one of them with a piece of food of the animal’s liking (same type of meat as in the training phase; baited cup) and adding five pieces of kibble (dry food) that produced a rattling sound when shaken, while the other cup remained empty (empty cup). At the beginning of each trial, the subject was lured away by throwing a piece of food far into the enclosure in line with the center of the table, to ensure that the animals approached the table centrally. This piece of food was usually a kibble pellet, but in some cases sausage or beef was used due to motivational reasons (this piece of food was, however, always considered of a “lower quality food” than the reward in the baited cup for each of the animals). As soon as the subject looked away, the experimenter placed the cups on the table, each one protruding out of each side of the curtain. Subsequently, the experimenter called the animal’s name or knocked in the middle of the table to draw their attention back to the apparatus.

The manipulation of the cups began as soon as the subject looked in the direction of the table (something that the experimenter could see on the video feed of the GoPro camera). Depending on the condition, one, both (one after the other) or no cups were manipulated by lifting and shaking them for several seconds until the subjects directed their attention towards them, with the experimenter placing them back on the table after doing so. At the end of the manipulation, the experimenter pushed the two targets through the fence to allow the subjects to make their choice by touching one of the targets with either their snout or their paws. After the animal touched one of the targets, both targets were pulled back to their original position behind the fence. If the subject chose the target on the side of the table where the baited cup was placed, the choice was considered correct. In this case, the reward was taken out of the cup within view of the subject and thrown over the fence into the enclosure. Conversely, if the subject chose the side where the empty cup was placed, the choice was considered incorrect. In this case, the cups were directly taken back behind the curtain and no reward was given. Only the first touch to the targets was considered for the purposes of the test and the subsequent data analyses, that is to say, if the animals managed to touch a second target before the experimenter pulled the targets out of reach of the animals, these additional touches were not taken into consideration.

The choice of the subjects was recorded by the experimenter on a prepared “protocol sheet”, and digitized after the session. This “protocol sheet” also contained the information about each of the trials the experimenter needed to conduct (i.e., the side on which the reward should be positioned and which cups (if any) should be shaken and in which sequence (in the case of the full condition).

For a video of a sample trial (full information condition), see **Video S1** in the supplementary materials.

##### Conditions

To examine the inferential reasoning abilities or potential alternative strategies the animals might have used in the study by Lampe et al. (2017), we confronted the animals with four different conditions (three experimental —full information, affirming the consequent, and denying the consequent— and one control condition —no information).

In the “full information” condition (similar to the one performed by Lampe and colleagues) both cups were shaken one after the other, with the baited cup emitting a rattling sound and the empty cup remaining silent. The order in which the cups were shaken in the “full information” condition (“baited” or “empty” first) was counterbalanced. Furthermore, in the “affirming the consequent” condition only the baited cup was shaken, emitting a rattling sound; while in the “denying the consequent” condition only the empty cup was shaken, emitting no sound. In the “no information” condition, no cup was shaken, and thus no information about the location of the reward was given to rule out possible alternatives strategies the animals might use to solve the task.

Within each test session, two trials per condition were presented in a pseudo-randomized pattern; in such a way that trials of the same condition never occurred more than twice in a row and the reward was not on the same side more than twice in a row.

A detailed overview of the conditions used in this experiment is shown in **Table 3**.

**Table 3:**
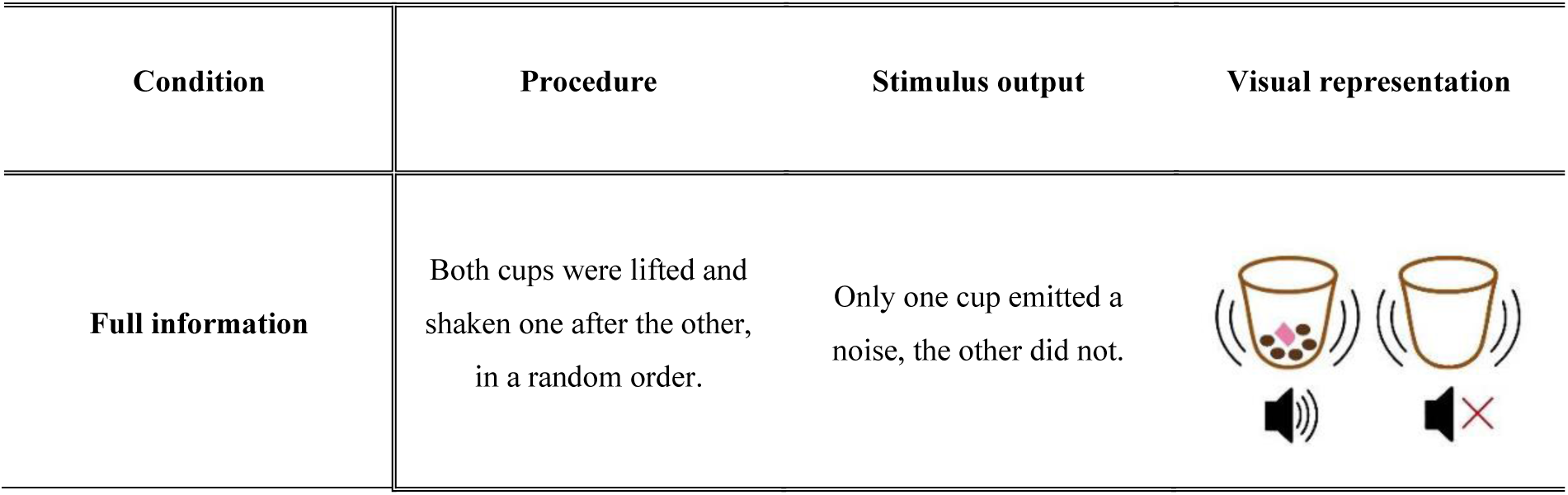
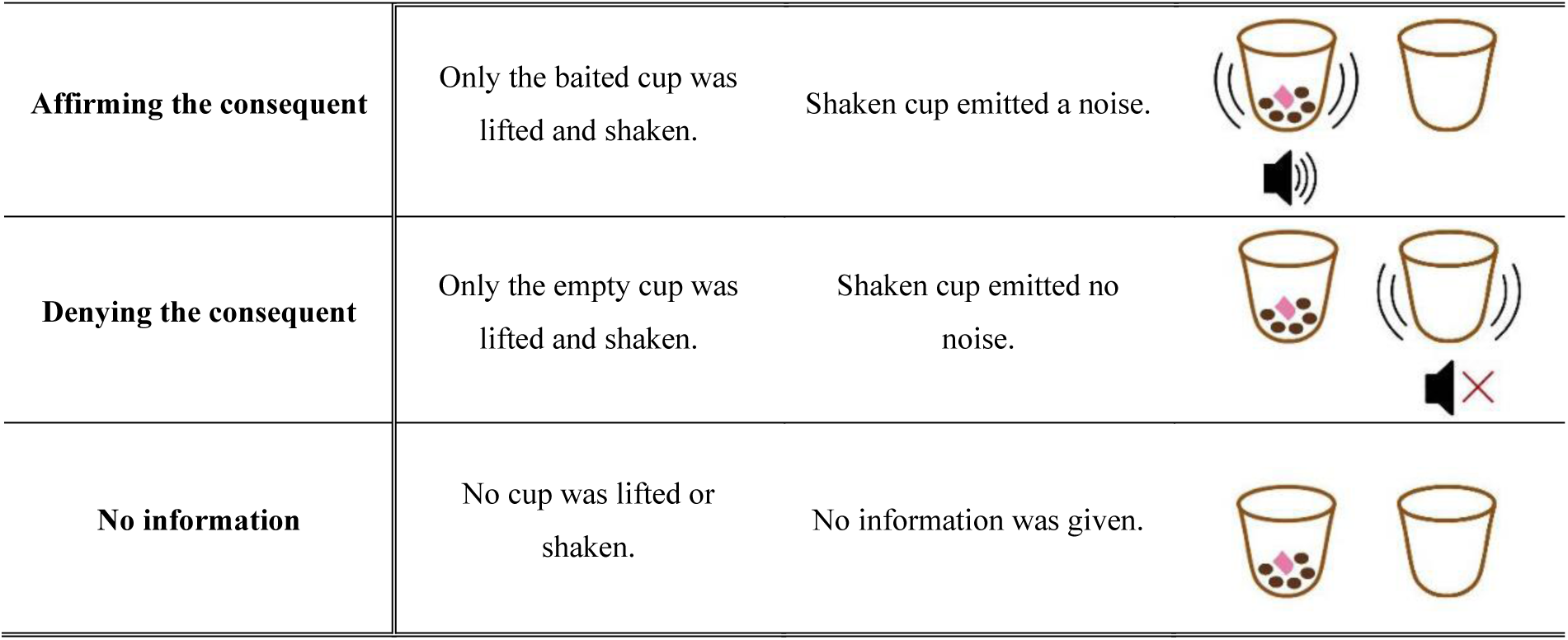
Description of the conditions used in this experiment. The pictures represent the cups (light brown) with one of them containing 5 kibble pellets (dark brown) and one piece of the meat-reward (pink). The kinetic lines indicate which cup is shaken in each condition, while the speaker icon represents the auditory information (rattling noise or silent) of the respective cup above it.

| Condition | Procedure | Stimulus output | Visual representation |
| --- | --- | --- | --- |
| Full information | Both cups were lifted and shaken one after the other, in a random order. | Only one cup emitted a noise, the other did not. |  |
| **Affirming the consequent** | Only the baited cup was lifted and shaken. | Shaken cup emitted a noise. |  |
| **Denying the consequent** | Only the empty cup was lifted and shaken. | Shaken cup emitted no noise. |  |
| **No information** | No cup was lifted or shaken. | No information was given. |  |

### Statistical analyses

The analyses were performed using R 4.0.3 (R Core Team 2020). To test if the performance of each species was above chance in each of the conditions, exact binomial tests (two-tailed) with a probability of success of 0.5 were used. To further examine the effects of the condition and the species on the experiment’s performance, we ran generalized linear mixed models (GLMM) using the “glmer” function (package “lme4”; Bates et al. 2015) with a binomial distribution and logit link function. Further, we checked for model assumptions through custom functions designed by Roger Mundry and Remco Folkertsma.

Our main model investigated whether wolves and dogs performed differently across the conditions. Thus, we used the outcome of each trial (“success” or “failure”) as the response variable, the species, the condition, and their interaction as predictor variables (fixed effects). We added the session in which each trial was performed as a control variable (fixed effect) to account for any variations in performance due to learning or changes in motivation throughout the experiment. Additionally, since the wolves were not fed every day (as opposed to the dogs) which might have caused difference in motivation, we also added the days that passed in between the date the animal was last fed and the day of the experiment (set as 0 if the subject was fed on the day before the test, 1 if one day had passed in between, and so on) as a control variable (fixed effect). The identity of the subjects was added as a random effect, and random slopes were added whenever applicable.

To test the effect of the interaction between the condition and the species, we performed a *χ*^2^ test using an “anova” (“car” package; Fox and Weisberg 2019) to compare the full model described above with a null model in which the variables species and condition were removed. To further investigate the effect of the species and the condition, we also ran reduced models that removed the interaction between these two factors, which were equally compared with the null model through a *χ*^2^ test.

Additionally, we conducted an analysis for the first two sessions only, to examine the animals’ initial performance in a manner comparable with the experiment done by Lampe et al. (2017). In that initial experiment, the condition our paradigm was based upon was performed in 4 trials (in 2 different sessions) interspersed in trials testing for understanding of another causal cues and several social ones. Both the predictor and the control variables were the same as the model described above (except for the session number, which was excluded due to the limited number of sessions analyzed, which could have negatively impacted the model’s stability). As was the case for the main model, we ran a full-null comparison with a model in which the predictor variables (species, condition, and the interaction between them) were removed. We also run a reduced model (without the interaction between species and condition) and compared it to the null model as well.

Furthermore, in order to account for any potential alternative strategies, the animals might have used to solve the task (e.g. choosing the most salient or last stimulus), we ran two more binomial tests (to compare performance against chance) and their respective models (to compare performance between the species). To check for a potential preference towards the last cup shaken (recency effect) we analyzed whether the animals chose the last cup shaken in the “full information” condition, regardless of whether it made a rattling noise or not. Additionally, in order to analyze the animals’ preference towards the most salient stimulus (independent of whether it was baited), we ran a model that investigated to which extent the animals chose the only shaken cup in the two partial information conditions (i.e., the side with the most salient stimulus).

Plots were generated using the “ggplot” function (package “tidyverse”; Wickham et al. 2019). Significance brackets in the plots were added through the “ggsignif” package (Ahlmann-Eltze & Patil, 2021) and colors in figure 3 were adjusted through the use of “ggnewscale” (Campitelli, 2023).

### Ethical statement

All procedures were approved by the “Ethics and Animal Welfare Committee at the University of Veterinary Medicine Vienna” with the ETK number “ETK-050/04/2021”. Participation in the experiment was voluntary for all animals. Throughout the testing sessions, the subjects had permanent access to water. Regarding the pet dogs, the owners’ consent was given in advance, and said owners were also informed beforehand about the procedure of the study. At any time, the animals had the possibility to retreat into the enclosure in case they were uncomfortable with the experiments.

## Results

Overall, to reach the testing phase, wolves needed an average of 2.9 ± 0.5 training sessions, while the pack dogs needed 3.6 ± 1.2 sessions and the pet dogs needed 2.8 ± 0.5 sessions. One of the pack dogs (*Layla*) needed 5 training sessions to move forward to the testing phase, but after failing three different pre-trials in two different sessions, she received an additional 3 training sessions (for a total of 8). Furthermore, one of the pet dogs (*Hakima*) also performed 2 additional training sessions since he was the pilot subject. Since this pilot was carried out more than 2 months before he was tested, we decided to give him an additional 2 training sessions. All these additional sessions were included in the above-mentioned averages. Details for the performance of each animal in the training sessions can be found in **Table S1**, in the supplementary materials.

### Comparing dog populations (pack dogs vs. pet dogs)

Pack and pet dogs showed similar performances in all conditions, as well as in the average of the choices for the second cup in the full information condition, and the average of choices for the shaken cup in the partial information conditions (**Table 4**). Accordingly, we found no significant difference between pack dogs and pet dogs neither across sessions (full-null comparison: *χ*^2^ = 5.054, *p* = 0.653) nor in the first two sessions (full-null comparison: *χ*^2^ = 5.010, *p* **=** 0.286). Further, we also found no effect of pack vs. pet dogs regarding whether they chose the cup shaken second in the “full information” condition (full-null comparison: *χ*^2^ = 0.293, *p* **=** 0.588) nor whether they chose the shaken cup in the partial information conditions (full-null comparison: *χ*^2^ = 2.347, *p* **=** 0.125). Since pack and pet dogs did not show any significant differences in their performance, we grouped pack dogs and pet dogs together for further analyses, as experience seemed to have no influence on the performance of the dogs in this study. Consequently, only the two species groups (wolves and dogs) will be compared hereafter.

**Table 4:** Average correct choices for each condition and choices meant to measure potential biases (± standard error) for both pack and pet dogs.

|  | Pack dogs | Pet dogs |
| --- | --- | --- |
| No information | $0.562 \pm 0.223$ | $0.447 \pm 0.204$ |
| Full information | $0.550 \pm 0.224$ | $0.521 \pm 0.205$ |
| Affirming the consequent | $0.700 \pm 0.206$ | $0.750 \pm 0.177$ |
| Denying the consequent | $0.325 \pm 0.210$ | $0.229 \pm 0.172$ |
| Choosing the last cup<br>(full information) | $0.575 \pm 0.222$ | $0.625 \pm 0.199$ |
| Choosing the shaken cup<br>(partial information conditions) | $0.688 \pm 0.208$ | $0.760 \pm 0.174$ |

### Comparing species (wolves vs. dogs) across all sessions (1-8)

#### All conditions: General performance

In all models, the “no information” condition (**Table 7**) was taken into the intercept, in order to compare the rest of the experimental conditions (“full information”, “affirming the consequent”, and “denying the consequent”) with the control. The dogs’ performance was taken into the intercept for all models as well.

##### Full model

Our analyses showed that the interaction between species and condition significantly influenced the performance of the animals (full-null comparison: *χ*^2^ = 53.585, p **<** 0.001). However, a significant difference between species was only found in the “denying the consequent” condition (GLMM_full_: estimate ± SE = 0.986 ± 0.320, p = 0.002; see **Table 5**; **Figure 2**) with dogs’ performance (0.273 ± 0.135 on average) being significantly worse than wolves’ (0.469 ± 0.158) and the latter being at chance level (see **Table 7**). In all other experimental conditions, no significant difference between species was found (**Table 5**; **Figure 2**).

**Figure 2:**
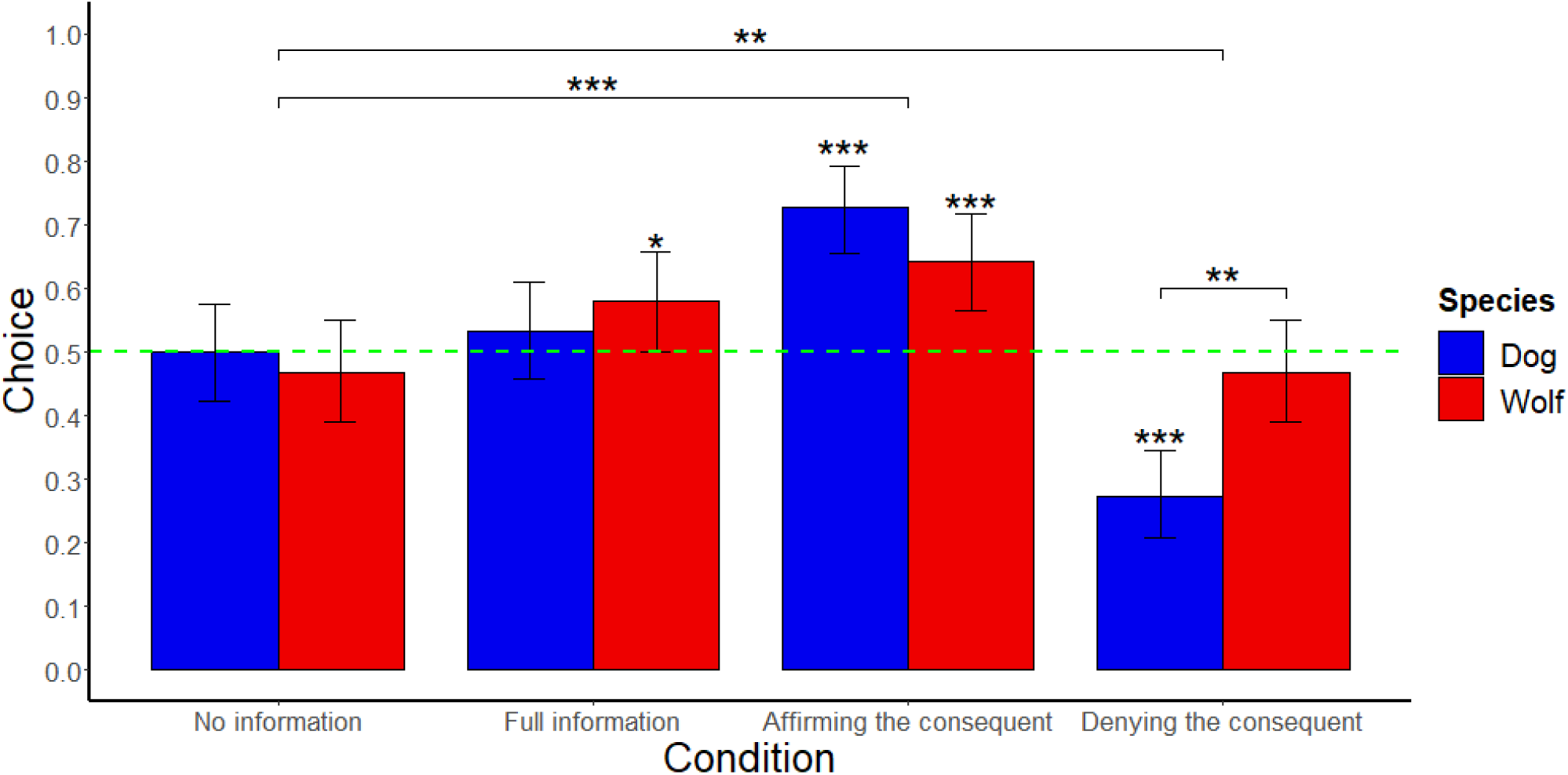
Performance of the two species (*N*_wolf_ = 10; *N*_dog_ = 11) in the respective conditions based on the mean success in all sessions. Error bars represent the 95% confidence interval. The green line indicates the probability of success at chance level. Performances above chance (two-tailed binomial test; probability of success = 0.5) are shown by black asterisks above the error bar, whereas performances below chance are shown by red asterisks above the error bar. Differences between species are based on the results of the full model (GLMM_full_) and are shown by a black line with asterisks above them. Differences between conditions are based on the results of the reduced model (GLMM_reduced_) and shown by a black line with asterisks above them. (‘*’: *p* < 0.05, ‘**’: *p* < 0.01, ‘***’: *p* < 0.001).

**Table 5:** Summary of the full model (GLMM_full_) examining the effect of the interaction between species and condition in all sessions. . The response variable (“success”) was defined as the choice of the noisy (baited) cup.

| Model terms | Estimate | SE | z-value | Pr(>\|z\|) |
| --- | --- | --- | --- | --- |
| (Intercept) | -0.019 | 0.158 | (-0.123) | (0.902) |
| Dog vs. Wolf | -0.083 | 0.237 | -0.352 | 0.725 |
| No information vs. full information | 0.137 | 0.214 | 0.642 | 0.521 |
| No information vs. affirming the consequent | 0.994 | 0.243 | 4.089 | < **0.001** |
| No information vs denying the consequent | -0.985 | 0.227 | -4.338 | < **0.001** |
| Session number (z-transformed) | -0.127 | 0.057 | -2.209 | **0.027** |
| Days passed between feeding and testing (z-transformed) | -0.035 | 0.073 | -0.481 | 0.630 |
| Dog vs. wolf: full information | 0.318 | 0.311 | 1.021 | 0.307 |
| Dog vs. wolf: affirming the consequent | -0.265 | 0.346 | -0.767 | 0.443 |
| Dog vs. wolf: denying the consequent | 0.986 | 0.320 | 3.084 | **0.002** |

Regarding the “full information” condition, although there was no significant difference between the species in the model (estimate ± SE = 0.318 ± 0.311, p = 0.307; see **Table 5**), wolves’ performance (0.581 ± 0.157 on average) was above chance level, while dogs’ performance (0.534 ± 0.151 on average) did not differ from chance (**Table 7**). It is important to note, however, that the lower boundary of the 95% confidence interval for the wolves’ “full information” condition was almost at the 50% “chance” level (0.501; **Table 7**), questionining whether the wolves did indeed performe above chance. In the “affirming the consequent” condition both wolves (0.644 ± 0.152 on average) and dogs (0.727 ± 0.135 on average) performed above chance level (**Table 6**) with no significant difference between them (GLMM_full_: estimate ± SE = -0.265 ± 0.346, p = 0.443; see **Table 5**).

**Table 6:** Summary of the reduced model (GLMM_reduced_) examining the effect of species and condition without the interaction of both in all sessions. . The response variable (“success”) was defined as the choice of the noisy (baited) cup. The predictors were species and condition, while the interaction between them was removed. As the “no information” condition was our control condition, it was defined as the baseline against which all experimental conditions were compared. Regarding species, dogs were defined as baseline against which wolves were compared. Significant values are shown in bold.

| Model terms | Estimate | SE | z-value | Pr(>\|z\|) |
| --- | --- | --- | --- | --- |
| (Intercept) | -0.143 | 0.132 | (-1.087) | (0.277) |
| Dog vs. wolf | 0.179 | 0.142 | 1.260 | 0.207 |
| No information vs. full information | 0.289 | 0.155 | 1.857 | 0.063 |
| No information vs. affirming the consequent | 0.872 | 0.189 | 4.610 | < **0.001** |
| No information vs. denying the consequent | -0.494 | 0.158 | -3.127 | **0.002** |
| Session number (z-transformed) | -0.125 | 0.057 | -2.194 | **0.028** |
| Days passed between feeding and testing (z-transformed) | -0.037 | 0.077 | -0.477 | 0.634 |

**Table 7:** Statistical values of binomial tests examining wolves’ and dogs’ performances under the respective conditions in all sessions. To test for performances above chance exact binomial tests (two-tailed) with a probability of success of 0.5 were used. Significant values are shown in bold representing performances that are significantly different from chance level.

| Condition | Species | Probability of success | 95% confidence interval | p-value |
| --- | --- | --- | --- | --- |
| No information | Wolf | 0.469 | 0.390 - 0.549 | 0.477 |
|  | Dog | 0.500 | 0.424 - 0.576 | 1.000 |
| Full information | Wolf | 0.581 | 0.501 - 0.659 | **0.048** |
|  | Dog | 0.534 | 0.457 - 0.609 | 0.407 |
| Affirming the consequent | Wolf | 0.644 | 0.564 - 0.718 | **< 0.001** |
|  | Dog | 0.727 | 0.655 - 0.792 | **< 0.001** |
| Denying the consequent | Wolf | 0.469 | 0.390 - 0.549 | 0.477 |
|  | Dog | 0.273 | 0.208 - 0.345 | **< 0.001** |

##### Reduced model

To test whether there were differences between the conditions in both species, and whether wolves and dogs had any general differences in performance, we ran a reduced model by removing the interaction between condition and species. The analysis showed that there was a significant difference between the reduced and the null model (reduced-null comparison: *χ*^2^ = 38.934, *p* < 0.001), with no effect of species (estimate ± SE = 0.179 ± 0.142, p = 0.207), but with all conditions either being significantly different from the no information condition or having a trend towards significance (see **Table 6**). Additionally, a negative effect of the session number was found (estimate ± SE = -0.125 ± 0.057, p = 0.028; **Table 6**; **Figure 3**), implying that performance gradually decreased as the experiment progressed.

**Figure 3:**
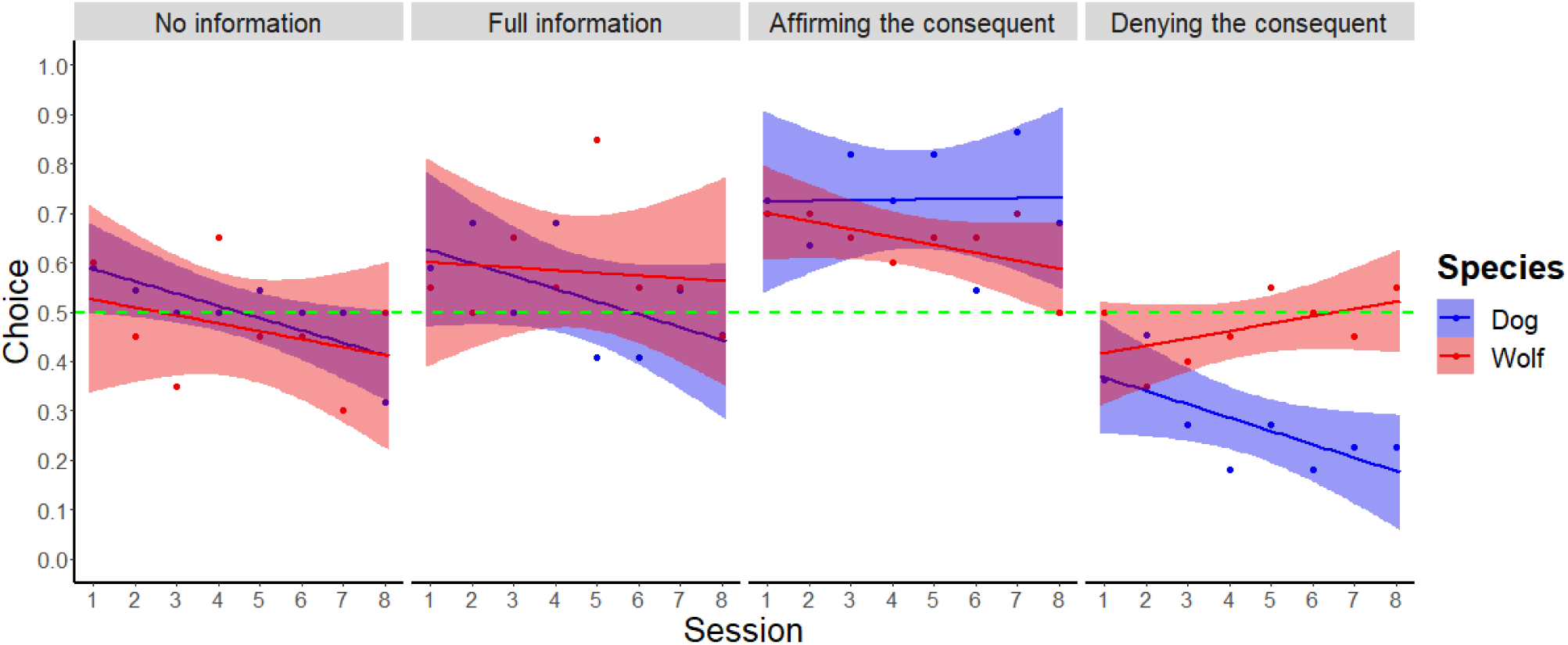
Performance over time of both species (*N*_wolf_ = 10; *N*_dog_ = 11) in each condition. Linear regression lines were applied by a linear model. The green line indicates the probability of success at chance level.

In terms of the effect of condition on performance, both partial information conditions (“affirming the consequent” and “denying the consequent”) differed significantly from the control condition (“no information”) (**Table 6**; **Figure 2**). In the “affirming the consequent” condition, subjects chose the correct side more often than in the “no information” condition (estimate ± SE = 0.872 ± 0.189, p < 0.001; **Table 6**). In contrast, in the “denying the consequent” condition subjects performed worse than in the “no information” condition (estimate ± SE = -0.494 ± 0.158, p = 0.002; **Table 6**).

Regarding the “full information” condition, we only found a trend (i.e., p-value greater than 0.05 but lesser than 0.10) in the comparison with the “no information” condition (estimate ± SE = 0.289 ± 0.155, p = 0.063) (**Table 6**; **Figure 2**).

The predictors were species, condition and their interaction. As the “no information” condition was our control condition, it was defined as the baseline against which all experimental conditions were compared. Regarding species, dogs were defined as baseline against which wolves were compared. Interaction of species and condition is indicated by “:”. Significant values are shown in bold.

##### Comparing species (wolves vs. dogs) and conditions in session 1 and 2

In the study done by Lampe et al. (2017), the animals were tested in a total of only four trials (two trials per session; two sessions). Accordingly, the version of the cups task done by Lampe et al. (2017) would be equivalent to the first four trials of the “full information” condition in ours. Consequently, in order to investigate the animals’ initial performance and compare the results from the “full information” condition with the study of Lampe et al. (2017), we decided to analyze the performance of the first two sessions separately.

No effect of the interaction between species and condition on the performance of the animals was found in this model (full-null comparison: *χ*^2^ = 12.094, *p* = 0.097). We also fit a second model in which the interaction was removed to test the effect on the performance of species and condition separately (GLMM_reduced_). We did find a significant difference between this model and the null (reduced-null comparison: *χ*^2^ = 11.157, *p* = 0.024); however, none of the experimental conditions differed significantly from the control (although there was a trend for the affirming the consequent condition; estimate ± SE = 0.613 ± 0.323, p = 0.057; see **Table 8** and **Figure 4** for more details).

**Figure 4:**
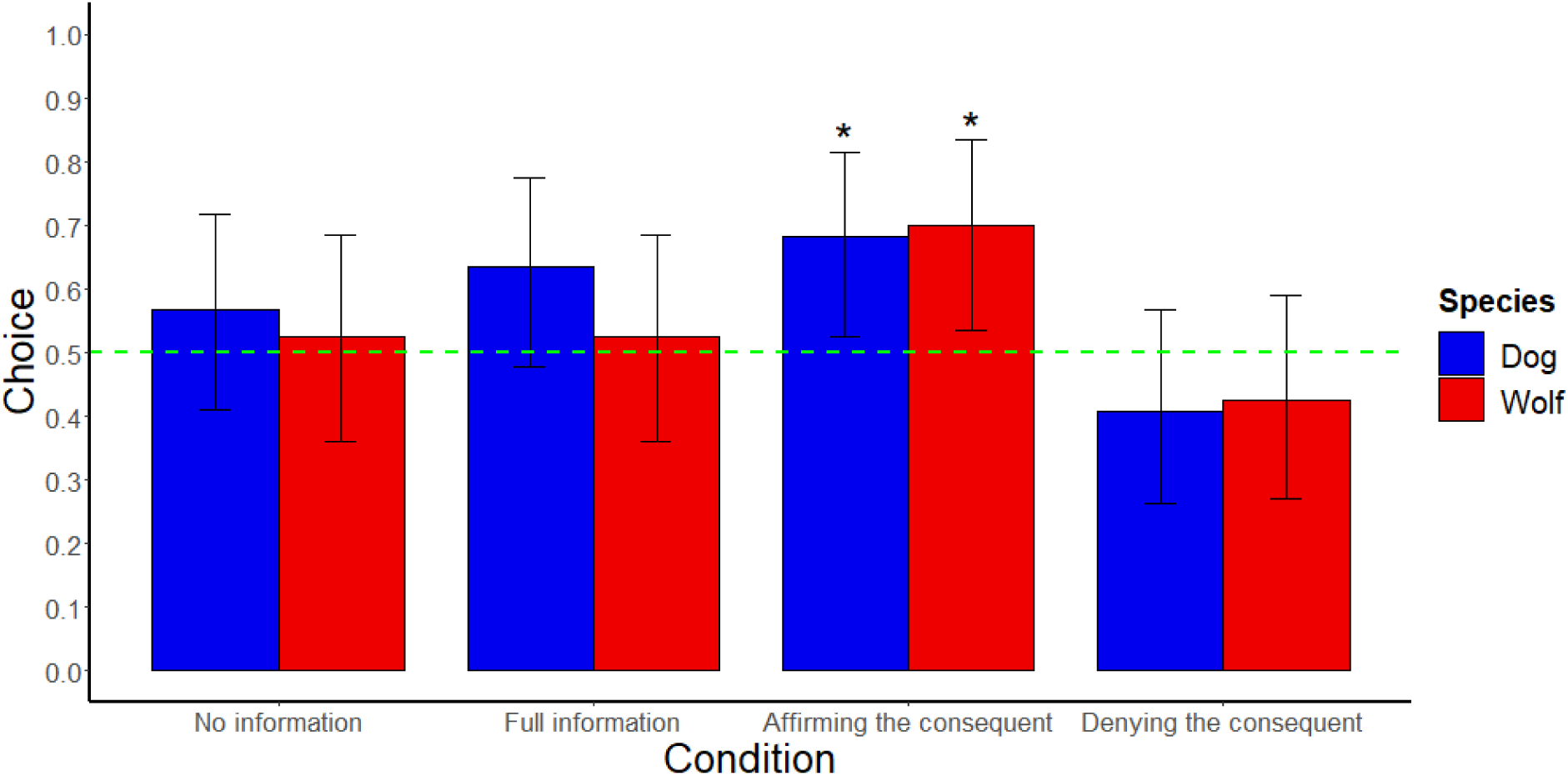
Performance of the two species (N_wolf_ = 10; N_dog_ = 11) in the respective conditions based on the mean success in the first two sessions. Error bars represent the 95% confidence interval. The red line indicates the probability of success at chance level. Performances above chance (two-tailed binomial test; probability of success = 0.5) are shown by black asterisks above the error bar. (‘*’: *p* < 0.05, ‘**’: *p* < 0.01, ‘***’: *p* < 0.001)

**Table 8:** Summary of the reduced model (GLMM_reduced_) examining the effect of species and condition without the interaction of both in the first two sessions. . The response variable (“success”) was defined as the choice of the noisy (baited) cup. The predictors were species and condition, while the interaction between them was removed. As the “no information” condition was our control condition, it was defined as the baseline against which all experimental conditions were compared. Regarding species, dogs were defined as baseline against which wolves were compared. Significant values are shown in bold.

| Model terms | Estimate | SE | z-value | Pr(>\|z\|) |
| --- | --- | --- | --- | --- |
| (Intercept) | 0.265 | 0.263 | (1.006) | (0.314) |
| Dog vs. wolf | -0.154 | 0.302 | -0.509 | 0.611 |
| No information vs. full information | 0.146 | 0.312 | 0.469 | 0.639 |
| No information vs. affirming the consequent | 0.613 | 0.323 | 1.901 | 0.057 |
| No information vs. denying the consequent | -0.535 | 0.325 | -1.647 | 0.100 |
| Days passed between feeding and testing (z-transformed) | 0.019 | 0.151 | 0.128 | 0.898 |

Further, we found no effect of species on the performance in the first two sessions, as the performance of the dogs did not differ significantly from that of the wolves (estimate ± SE = -0.154 ± 0.302, p = 0.611; see **Table 8**). Both species performed at chance level in the “no information” condition, the “full information” condition and the “denying the consequent” condition (**Table 9**). Only in the “affirming the consequent” condition wolves (0.700 ± 0.147) and dogs (0.681 ± 0.142) performed above chance level **(Table 9**; **Figure 4**).

**Table 9:** Statistical values of binomial tests examining wolves’ and dogs’ performances under the respective conditions in the first two sessions. To test for performances above chance exact binomial tests (two-tailed) with a probability of success of 0.5 were used. Significant values are shown in bold representing performances that are significantly different from chance level.

| Condition | Species | Probability of success | 95% confidence interval | p-value |
| --- | --- | --- | --- | --- |
| No information | Wolf | 0.525 | 0.361 - 0.685 | 0.875 |
|  | Dog | 0.568 | 0.410 - 0.717 | 0.451 |
| Full information | Wolf | 0.525 | 0.361 - 0.685 | 0.875 |
|  | Dog | 0.636 | 0.477 - 0.776 | 0.096 |
| Affirming the consequent | Wolf | 0.700 | 0.535 - 0.834 | **0.017** |
|  | Dog | 0.682 | 0.524 - 0.814 | **0.023** |
| Denying the consequent | Wolf | 0.425 | 0.270 - 0.591 | 0.430 |
|  | Dog | 0.409 | 0.263 - 0.568 | 0.291 |

##### Bias towards the second cup shaken in the “full information condition”

Based on the performance in the full information condition, it is not clear whether the animals used inferential reasoning, followed the most salient stimulus, or chose the last stimulus presented. Therefore, in an additional analysis, the choice of the last cup shaken in the “full information” condition was examined.

The results show that both wolves and dogs chose the cup shaken last in the “full information” condition above chance level (binomial test: Wolves: probability of success: 0.656, C.I. 95%: 0.577 - 0.729, *p* < 0.001; Dogs: probability of success: 0.602, C.I. 95%: 0.526 - 0.675, *p* = 0.017; see **Figure 5**). This suggests that both species’ choices were at least partially informed by the position of the last cup shaken (recency effect).

**Figure 5:**
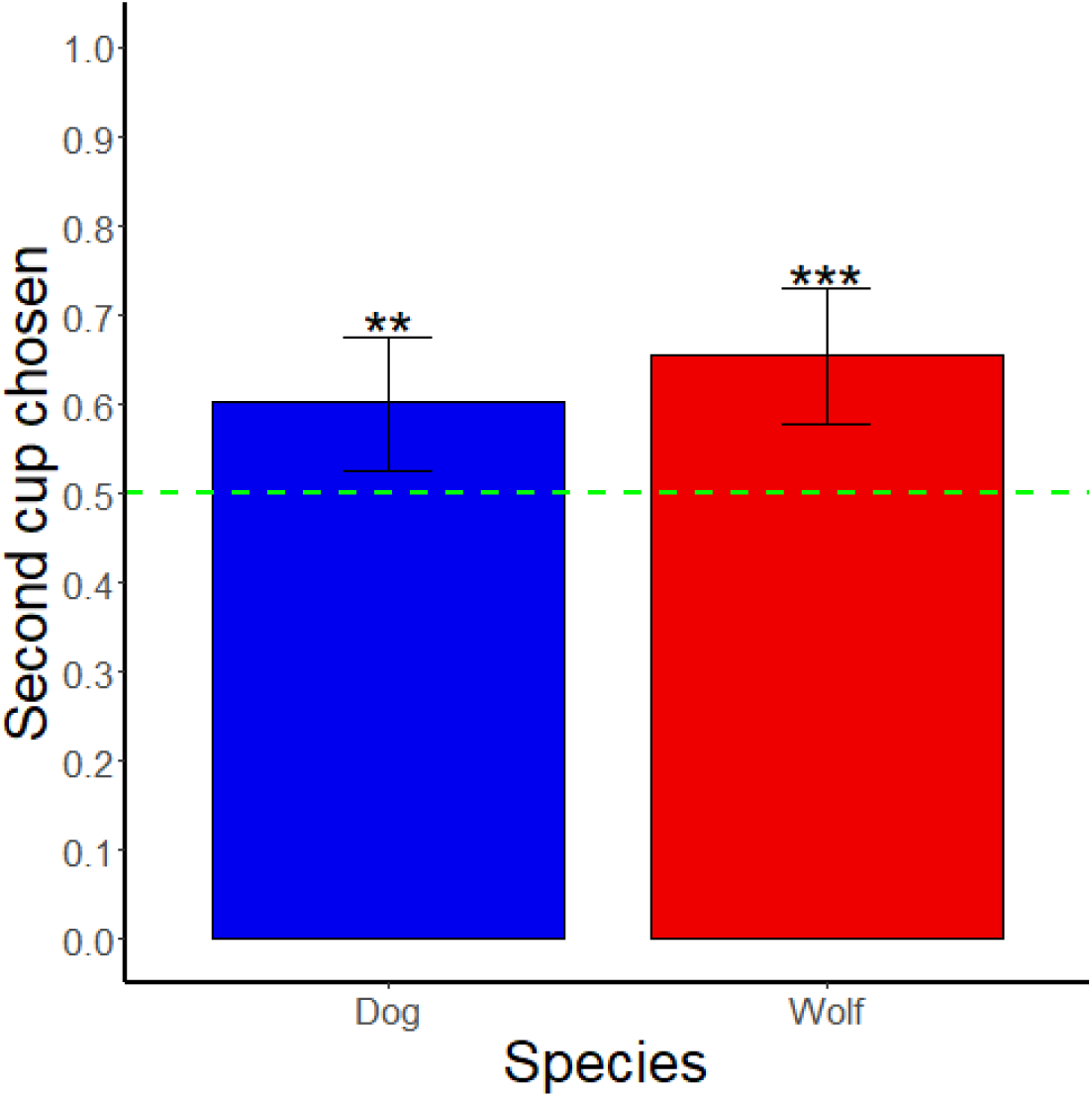
Performance of the two species (N_wolf_ = 10; N_dog_ = 11) regarding the choice of the second shaken cup in the full information” condition. Error bars represent the 95% confidence interval. The green line indicates the probability of success at chance level. Performances above chance (two-tailed binomial test; probability of success = 0.5) are shown by black asterisks above the error bar. (‘*’: *p* < 0.05, ‘**’: *p* < 0.01, ‘***’: *p* < 0.001).

Additionally, to test whether there were differences between the species in this preference towards the second cup shaken, we performed a *χ*^2^ test comparing the full model that included species as a predictor variable with a null model without this factor. No significant difference between the models was found (full-null comparison: *χ^2^*= 1.198*, p* = 0.274; see **Figure 5**), indicating that there were no differences between them in their bias towards the second/last cup.

However, considering the individual performance of the animals, the results show large variation between subjects in both species regarding the choice of the cup shaken second when it was baited or not (see **Figure S1** in the supplementary material). For example, three wolves (*Taima*, *Amarok*, and *Etu*) and three dogs (*Imara*, *Asali*, and *Zazu*) chose the cup shaken second in 12 or more out of 16 trials (see **Figure S2** in the supplementary material). Only one wolf (*Chitto*) chose the baited cup in the full information condition in a total of 12 trials (6 trials baited cup shaken first; 6 trials baited cup shaken second), and only one dog (*Hakima*) in a total of 11 trials (4 trials baited cup shaken first; 7 trials baited cup shaken second), while no other subject achieved these performances in the “full information” condition (see **Figure S1** in the supplementary material).

##### Bias towards the most salient stimulus in the partial information conditions

To further investigate which strategies the animals used, we also analyzed whether the animals had a preference towards the most salient stimulus (i.e. the cup that was shaken) in the partial information conditions (“affirming the consequent” and “denying the consequent”).

When taking the data from the two partial information conditions, both species chose the cup that was shaken (saliency effect), regardless of whether it was the baited or empty cup, above chance level (binomial test: Wolves: probability of success: 0.588, C.I. 95%: 0.531 - 0.642, *p* = 0.002; Dogs: probability of success: 0.727, C.I. 95%: 0.678 - 0.773, *p* < 0.001; see **Figure 6**). Further, we found an effect of species (full-null comparison: *χ*^2^ = 7.401, *p* = 0.007) with a significant difference between wolves and dogs (GLMM_full_: estimate ± SE = -0.566 ± 0.206, *p* = 0.006); with dogs choosing the cup that was shaken (most salient stimulus) significantly more often than wolves (**Figure 6**; see also **Table S2**).

**Figure 6:**
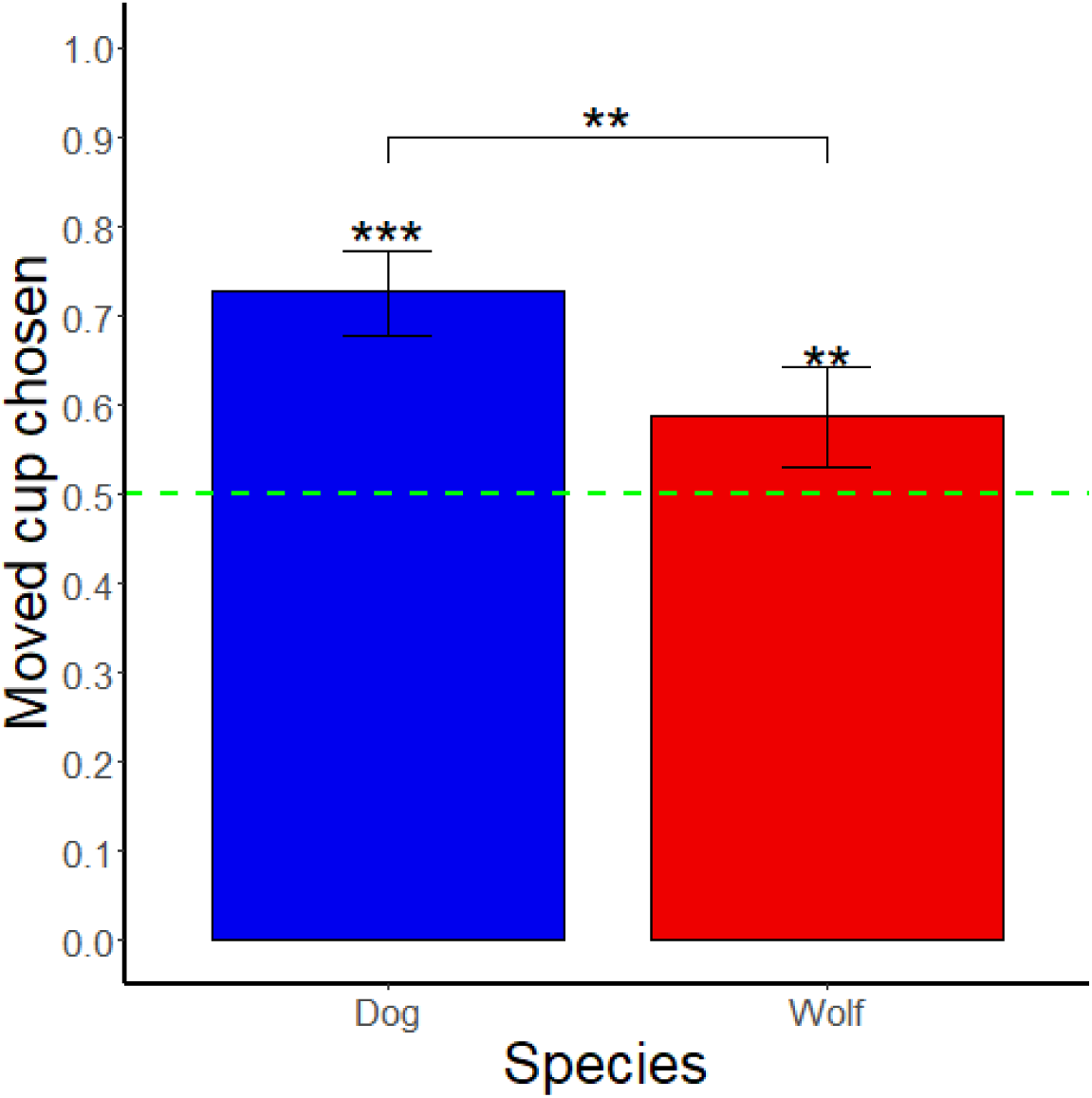
Performance of the two species (*N*_wolf_ = 10; *N*_dog_ = 11) regarding the choice of the only moved cup in the partial information conditions. Error bars represent the 95% confidence interval. The green line indicates the probability of success at chance level. Performances above chance (two-tailed binomial test; probability of success = 0.5) are shown by black asterisks above the error bar. Differences between species are based on the results of the full model (GLMM_full_) and are shown by a black line with asterisks above them. (‘*’: *p* < 0.05, ‘**’: *p* < 0.01, ‘***’: *p* < 0.001).

## Discussion

In this study, we investigated whether wolves and dogs (pack and pet dogs) use inferential reasoning to infer the location of a hidden reward by using the presence or absence of sound to inform their choices. Our results did not reveal any significant differences between pet dogs and pack dogs, thus we pooled the data. We found that wolves and dogs solved the “affirming the consequent” (only baited cup shaken) condition but not the “full information” condition (both cups shaken). Furthermore, dogs performed below chance levels in the “denying the consequent” condition (only empty cup shaken). Both species had a preference towards the last cup shaken in the full information condition, and towards the only cup shaken in the partial information conditions.

Considering our results, the strategy that was most likely employed by both species does not necessarily involve the use of inferential reasoning, but rather relies on the animals responding to other simple cues available. Judging by the results, the choices made by the wolves and the dogs responded both to a recency effect (bias towards the last stimulus presented) and a saliency effect (bias towards the most conspicuous stimulus presented), with the latter being more pronounced in the case of the dogs.

The effect of salience is well-known for influencing associative learning processes, with the stimuli that are proportionally more intense (in comparison with other stimuli) favoring faster associations (Frieman & Reilly, 2015). This effect has been suspected to interfere with a multitude of experiments in the field of animal behavior, with one stimulus or feature of the paradigm attracting the attention of the subjects regardless of whether it was conducive to solve the task (see Eckert et al., 2022; Lazzaroni et al., 2020; Marshall-Pescini et al., 2012 for a few examples). In regard to studies examining inference in animals, the possibility that a different stimulus might overshadow their potential inferential reasoning abilities has been considered several times in the past (see Mikolasch et al., 2012; O’Hara et al., 2015; O’Hara et al., 2016; Shaw et al., 2013 for a few examples). In our experiment, subjects seemed to pay attention to the cups’ movement above any other features of it, something that matches the results of previous studies on inference in dogs (Bräuer et al., 2006; Erdőhegyi et al., 2007). This could potentially respond to these species’ prey drive, to their history of training and participation with other tasks that imply movement, or both.

Recency effects are equally well-known, and have been studied in both humans and non-human animals across a range of cognitive tasks (Castro & Larsen, 1992; Murphy et al., 2006; Rubio et al., 2023, Thomas et al., 1984). In dogs, it has been shown that they place a higher value upon the most recently-acquired information when it comes to foraging (Devenport & Devenport, 1993). Accordingly, the movement of the first cup could have been “de-valued” by the time the second cup was presented. This issue could theoretically be tackled in future versions of this paradigm by presenting both stimuli simultaneously. Unfortunately, due to physical limitations of our apparatus (chiefly amongst which, the fact that the targets needed to be far enough from each other to avoid cross-associations with the wrong target), presenting both cups simultaneously could risk the animals failing to pay attention to both of them at the same time.

Taken together, both wolves and dogs seemed to respond to the saliency of the stimuli and the order in which they were presented rather than performing any inferences about the position of the reward. Using these strategies is, while not optimal, also not disadvantageous. From the way our sessions were presented, choosing the most salient stimulus (which, judging by our results, would be the last cup that was moved) would yield a reward half of the time (as there was an equal number of trials for all four conditions: P_full information_ = 0.5, P_affirming_ = 1, P_denying_ = 0, P_no info_ = 0.5; P_total_ = 0.5), which may have been good enough for the animals to lack the need towards developing a better strategy.

Another strategy we encountered is side bias, as several individuals participating in this study (5 out of 10 wolves, 4 out of 5 pack dogs, and 2 out of 6 pet dogs) showed a tendency towards choosing one side of the apparatus (see Table S3 for more information). Indeed, bias towards one side when two options are presented to the animals is common in experiments involving non-human animals especially those involving complex paradigms (see Hare & Tomasello, 1999; Laschober et al., 2021; Szabo et al., 2019 for a few examples; see Bolló et al., 2023 for a study investigating the potential mechanisms behind side bias in dogs). Though research on the proximate causes of side biases remains scant, one of the potential reasons behind it is that the animals engage with the tasks in which this bias presents itself as if they were operant conditioning tasks with a variable ratio schedule of reinforcement (i.e., a reinforcement scheme in which rewards are handed out with a certain probability, but with no way of knowing which of the responses will be reinforced; Miltenberger, 2016). Thus, if the animal ignores the stimuli presented and engages solely (or most often) with one of the options, this behavior would be strongly reinforced, as rewards would be provided half of the time but at random intervals from the animal’s perspective (a process that has been described as the reason for the addictiveness of gambling; Lozano Bleda & Pérez Nieto, 2012).

Overall, our results seem to be consistent with the available literature for the “cups task” in dogs (Bräuer et al., 2006; Lampe et al., 2017), but not so for the one in wolves (Lampe et al., 2017). Indeed, in our study, wolves did not show a better performance in the full information condition than in the control, even though they did solve the task above chance in the study by Lampe and colleagues. Even when we analyzed the performance of the animals only in the first four trials of our “full information” condition to make our study more comparable with Lampe et al., wolves did not perform above chance in our experiment, nor were there any differences between the species. These differences could be due to two main procedural differences: First, Lampe and colleagues tested the animals in two different causal conditions (“shape”: two wooden shapes lying on the table, one hiding food underneath; “noise”: two shaken containers, one of which was baited and emitted a noise when shaken) as well as in several social conditions, that involved witnessing human actions. Thus, it is possible that these social conditions influenced the animals’ performance by increasing the animals’ attention towards the experiment. In contrast, our study did not include any social conditions, which might have reduced the animals’ motivation. In fact, we observed a decrease in performance throughout our experiment, which could be explained by losing motivation towards carrying out the experiment. Second, the two cups were always shaken in the same order in the “noise” condition in Lampe’s study: first the baited cup, and then the empty cup. Considering that the most salient stimulus was presented always first in Lampe and colleagues’ study, saliency may have played an even larger role in that study than in ours, as going towards the most salient stimulus and ignoring any other, less salient, stimulus presented afterwards could explain the results observed and would have been the optimal strategy in this scenario. The recency effect in our study could have been due to the increased complexity as a result of 1) the counterbalancing, as in half of the trials the animals had to choose the first object and ignore the second, while the opposite held true in the other half and/or 2) presenting too many trials that, although different in function, may have looked too similar to each other in practice, which may have confused the animals.

Interestingly, while at first glance, dogs’ performance in the previous study was at chance level, dogs seemed to have tended to select the most recent stimulus (percentage of trials in which the dogs chose the last stimulus: pack dogs: 0.61 ± 0.14; pet dogs: 0.59 ± 0.16). This statistical non-significance could be explained by dogs having received on average less trials than wolves (wolves: 4 trials for all animals, pack dogs 3.17 ± 0.30 trials per animal, pet dogs 3.20 ± 0.33 trials per animal), allowing for the possibility of dogs showing a significant recency effect in Lampe and colleagues’ study as well if they were tested in 4 trials. Alternatively, dogs’ performance in Lampe et al.’s study could also be explained by saliency alone, assuming that the most salient feature for them was the movement in and of itself rather than the combination of movement and noise (as it seemed to be the case in our experiment).

Taken together, both our results and those presented by Lampe and colleagues can be explained through the effect of saliency (and to a lesser extent, that of recency) alone, albeit these effects did not show in quite the same manner between species and studies.

To put all of this within the wider context of the application of the auditory version of the cups task across species, wolves and dogs’ performance seems to somewhat relate to that of non-ape animals (and even some ape species, with orangutans showing only partial preference towards the container that made a sound or no preference at all; Call, 2004; De Petrillo & Rosati, 2020; Hill et al., 2011). Squirrel monkeys (Marsh et al., 2015), for instance, solved the task above chance when only the baited cup was shaken, did so as well when both the baited and empty cups were shaken (although one of the three subjects failed to do so), and chose below chance when only the empty cup was shaken; a performance somewhat resembling what we observed in our study (and equally telling of a potential saliency effect).

Comparisons between studies following this paradigm are, however, rather difficult to make in general, as different versions of it may change the testing regiment in small but crucial ways. For example, in an experiment done in African grey parrots (Schloegl et al., 2012), the subjects that were not successful in both of the equivalents to our full information and affirming the consequent conditions underwent a training phase before being tested again. In this training, they were shown the inside of the empty cup and the fact that it made no noise when shaken, and also that dropping a piece of food inside the cup and shaking it afterwards would make a noise. Similar to this, another experiment done with capuchin monkeys (Sabbatini & Visalberghi, 2008) allowed the animals to freely interact with the containers after finding that only a few animals were able to solve the task. They were re-tested after this exposure, to a somewhat higher rate of success (1 out of 8 animals solving the task in the first testing phase versus 3 out of 8 in the one after the exposure). As opposed to both of these studies, we did not use any of these training methods, as our subjects never saw the contents of the cups throughout the experiment, and no other demonstration that could convey that a baited cup would make noise and empty cup would make none was provided either.

The question arises, then, as to what prior knowledge of the contingencies should be expected of the animals attempting to solve this task. From a modern human’s *umwelt* (*sensu* Uexküll & Mackinnon, 1926), a noise coming from a moving opaque container would of course signify the presence of something inside it. However, most animals do lack any prior experience with this particular scenario (and thus, they are not equipped to interact with the task in ways beyond the saliency of the stimulus at hand). All of our subjects had prior experience with the sound of kibble pellets against plastic due to their daily feeding and other experiments (and indeed, some of them had been previously tested in Lampe et al.’s version of the experiment), but none had experience with the specific features of this version of the experiment. Therefore, though it could be argued that a capacity to make inferences based on auditory stimuli —potentially used in the wild by predators to locate their prey— could be employed to solve a task such as this, it is unclear whether such an ability would require prior experience with the stimulus used in order to make inferences about it.

In order to tackle the issues intrinsic to this paradigm (and in particular, to the current study’s implementation of it), we propose future endeavors regarding the study of inferential reasoning in wolves and dogs to focus on the use of touchscreens and similar automatized apparatuses. Touchscreen-based apparatuses have been successfully used in dogs and wolves in the past (e.g., Aust et al., 2008; Dale et al., 2019; Laude et al., 2016; Range et al., 2008; Rivas-Blanco et al., 2020), and indeed, they have also been successfully used to investigate inferential reasoning capabilities in other species (Aust et al., 2008; Mikolasch et al., 2013; O’Hara et al., 2015; O’Hara et al., 2016), as well as dogs themselves (Aust et al., 2008). The stimulus presented through such means would be simpler and easier to control, thus reducing the possibility of the animals making associations towards irrelevant features of the task, and indeed tackling the question of how much previous knowledge on the contingencies of the task would be required for it to be solved (as all the animals would need to be trained on the stimuli presented). Furthermore, by removing the need for an experimenter, any potential distractions and cueing derived from their presence would be removed as well. Additionally, an apparatus such as this could either employ motion sensors to start each trial only when the subject is positioned at a certain distance of the apparatus or start the trials by the animal’s contact with the apparatus, thus eliminating the need to use food to reposition them between trials (and thus, the potentially confounding factor of being rewarded even when the *incorrect* choice was made).

In conclusion, we found neither sufficient evidence that wolves employ causal reasoning nor any differences in performance between wolves and dogs. Instead, our results show that both wolves and dogs favored both the most salient stimulus and the most recent one presented, potentially due to confounding factors that arose from the manner in which the paradigm was implemented. Thus, it remains unclear whether the animals lack the ability for inferential reasoning or rather it was the experimental design of the current study, which caused unintended distractions that biased the animals’ performance. Future studies should consider the use of automatized apparatuses in order to reduce these potential distractions.

## Supporting information

Supplementary video (set up)

## Acknowledgements

We thank Kathryn Roddie and the trainers at the Wolf Science Center Core facility for contributing to the data collection process. We also thank Erik Hollerweger and Kathryn Roddie for crafting the apparatus used for this experiment.

## Author contributions

FR and SMP conceptualized the experiment; DRB developed the methodology; SK and DRB performed the data collection; DRB and SK curated and analyzed the data; SK, DRB, and FR wrote the first draft of the paper; FR and SMP supervised the entirety of the experiment; all authors edited and reviewed the manuscript.

## Funding

This study was funded by the Austrian Science Fund (Fonds zur Förderung der wissenschaftlichen Forschung, FWF), FWF grant number P33928-B. DRB was funded by a Marietta Blau Grant from the Austrian Agency for International Cooperation in Education, Science and Research (MMC-2023-07030).

## Data avaliability

All data generated or analyzed during this study are included in this published article as supplementary materials.

## Notes

### Competing Interest Statement

The authors have declared no competing interest.

