## Supplementary material (tables and figures) for "Inference in wolves and dogs: The “cups task”, revisited"

### Supplementary materials

**Table S1: List of animals with additional information regarding their exact date of birth, breed, date when they started the first training session and testing enclosure**. The genetic relationship is represented by letters: animals from the same litter will share a letter, and whenever the letter is accompanied of a number (a "1" in all cases), that subject was the offspring from a member of the litter represented by that letter. The horizontal division lines indicate the packs or households the animals live in together. Animals shown in grey and italics were excluded from the experiment due to motivational or health problems. Animals presented in bold were former pack dogs which are living in human families for several years.

| Species | Population | Subject | Sex | Litter | Birth date | Breed / subspecies | Start of experiment | Test enclosure | Training sessions to criterion |
|---|---|---|---|---|---|---|---|---|---|
| Wolf | pack | Chitto | m | A | 07/04/2012 | Timber wolf | 14/10/2021 | NTE | 2 |
| *Wolf* | *pack* | *Tala* | *f* | *B* | *04/04/2012* | *Timber wolf* | *14/10/2021* | *NTE* | — |
| *Wolf* | *pack* | *Nanuk* | *m* | *C* | *28/04/2009* | *Timber wolf* | *06/11/2021* | *9a* | — |
| Wolf | pack | Una | f | A | 07/04/2012 | Timber wolf | 07/11/2021 | 9a | 3 |
| Wolf | pack | Taima | f | D | 04/05/2016 | Timber-European wolf | 22/11/2021 | NTE | 7 |
| Wolf | pack | Tekoa | m | D | 04/05/2016 | Timber-European wolf | 15/11/2021 | NTE | 2 |
| Wolf | pack | Geronimo | m | E | 02/05/2009 | Timber wolf | 22/11/2021 | NTE | 2 |
| Wolf | pack | Yukon | f | E | 02/05/2009 | Timber wolf | 22/11/2021 | NTE | 2 |
| Wolf | pack | Etu | m | G | 04/05/2016 | Timber wolf | 06/04/2022 | 9a | 2 |
| Wolf | pack | Maikan | m | D | 04/05/2016 | Timber-European wolf | 11/04/2022 | 9a | 2 |
| Wolf | pack | Kenai | m | H | 01/04/2010 | Timber Wolf | 12/03/2022 | 9a | 4 |
| Wolf | pack | Amarok | m | B | 04/04/2012 | Timber Wolf | 12/03/2022 | 9a | 3 |
| Dog | pack | Hiari | m | I | 21/03/2014 | Mongrel | 05/12/2021 | NTE | 4 |
| Dog | pack | Imara | f | I | 21/03/2014 | Mongrel | 05/12/2021 | NTE | 2 |
| Dog | pack | Layla | f | J | 02/08/2011 | Mongrel | 06/12/2021 | NTE | 8(5+3)[1] |
| Dog | pack | Panya | f | J1 | 02/04/2014 | Mongrel | 06/12/2021 | NTE | 2 |
| Dog | pack | Enzi | m | J1 | 02/04/2014 | Mongrel | 12/12/2021 | NTE | 2 |
| Dog | pet | **Asali** | m | K | 19/09/2010 | Mongrel | 16/01/2022 | OTE | 2 |
| *Dog* | *pet* | ***Kilio*** | *m* | *L* | *18/12/2009* | *Mongrel* | *16/01/2022* | *OTE* | — |
| Dog | pet | **Hakima** | m | M | 19/09/2010 | Mongrel | 24/01/2022 | OTE | 4(2+2)[2] |
| Dog | pet | **Pepeo** | m | J1 | 02/04/2014 | Mongrel | 16/01/2022 | OTE | 2 |
| Dog | pet | Freya | f | N | 07/05/2015 | Terrier | 16/01/2022 | OTE | 2 |

[1] Failed 3 pre-trials after completing training (on her 5$^{th}$ session) and therefore received an additional 3 training sessions (for a total of 8)
[2] Received first 2 training sessions as a pilot subject and 2 more as a test subject, two months later (for a total of 4)

| | | | | | | | | | |
|---|---|---|---|---|---|---|---|---|---|
| Dog | pet | **Zazu** | m | J1 | 02/04/2014 | Mongrel | 17/01/2022 | OTE | 5 |
| Dog | pet | Coco | f | O | 21/02/2017 | Border Collie | 25/01/2022 | OTE | 2 |

**Figure S1: Individual performance of the animals in the "full information" condition**. The numbers represent the total number of trials chosen in each case: *dark green*: baited cup was shaken first it was chosen by the animal; *dark grey*: baited cup was shaken first, but the empty cup was chosen; *light green*: baited cup was shaken second and it was chosen by the animals; *light grey*: baited cup was shaken second, but empty cup was chosen. The black vertical line separates the species (left: wolves; right: dogs).

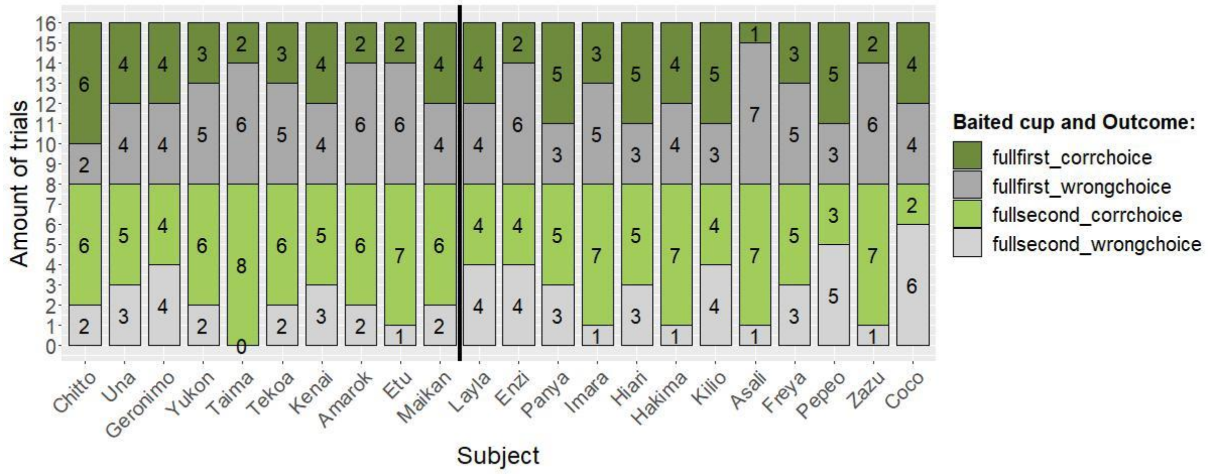


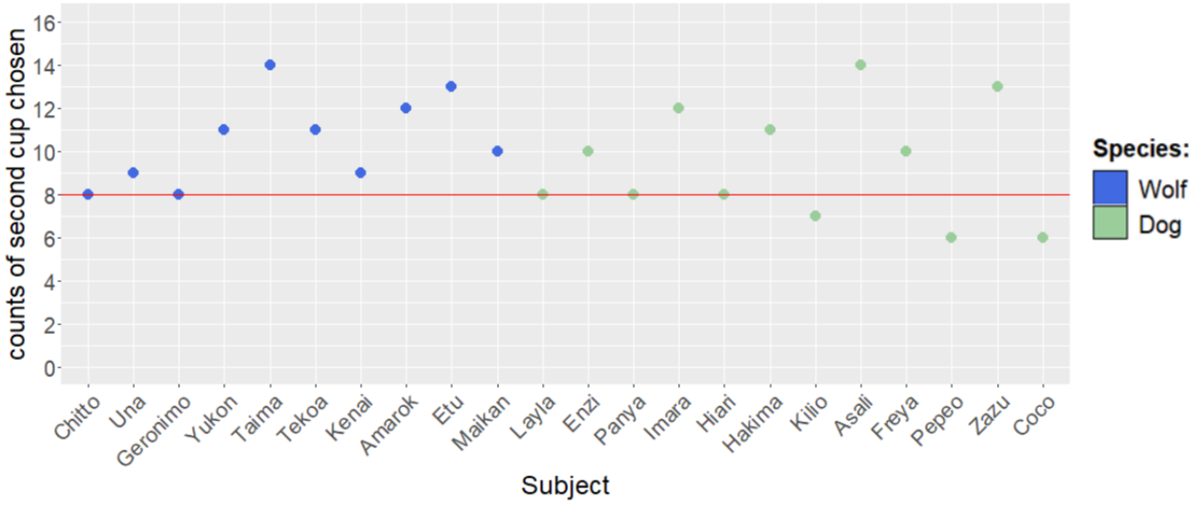


**Figure S2: Number of trials in which the second shaken cup was chosen in the "full information" condition, regardless of the position of the reward**. The red line indicates the probability at chance level.

**Table S2: Summary of the model (GLMM$_{full}$) examining the effect of species on the "moved cup" choice in the two partial information conditions**. Species was defined as the predictor variable, with dogs were defined as baseline against which wolves were compared. Significant values are shown in bold.

| **Model terms** | **Estimate** | **SE** | ***z*-value** | **Pr(>\|*z*\|)** |
|---|---|---|---|---|

| | | | | |
|---|---|---|---|---|
| dogs vs. wolves | -0.541 | 0.200 | -2.709 | **0.007** |
| session number (z-transformed) | 0.036 | 0.097 | 0.372 | 0.710 |
| days passed between feeding and testing (z-transformed) | -0.056 | 0.098 | -0.569 | 0.569 |

**Table S3: Statistical values of binomial tests examining subjects' individual tendencies to choose one side (arbitrarily set to be the left one).** To test for side biases, exact binomial tests (two-tailed) with a probability of success of 0.5 were used. Significant values are shown in bold representing performances that are significantly different from chance level (i.e., whenever subjects show to be biased towards either choosing the left or right side).

| **Subject** | **Population** | **Probability of choosing left** | **95% conficence interval** | **p-value** |
|---|---|---|---|---|
| Chitto | wolf | 0.656 | 0.527 — 0.771 | **0.017** |
| Una | wolf | 0.281 | 0.176 — 0.408 | **0.001** |
| Geronimo | wolf | 0.406 | 0.285 — 0.536 | 0.169 |
| Yukon | wolf | 0.391 | 0.271 — 0.521 | 0.103 |
| Taima | wolf | 0.484 | 0.358 — 0.613 | 0.901 |
| Tekoa | wolf | 0.500 | 0.372 — 0.628 | 1.000 |
| Maikan | wolf | 0.297 | 0.189 — 0.424 | **0.002** |
| Etu | wolf | 0.734 | 0.609 — 0.837 | **< 0.001** |
| Amarok | wolf | 0.453 | 0.328 — 0.583 | 0.532 |
| Kenai | wolf | 0.641 | 0.511 — 0.757 | **0.033** |
| Layla | pack dog | 0.781 | 0.660 — 0.875 | **< 0.001** |
| Enzi | pack dog | 0.797 | 0.678 — 0.887 | **< 0.001** |
| Panya | pack dog | 0.734 | 0.609 — 0.837 | **< 0.001** |
| Imara | pack dog | 0.578 | 0.448 — 0.701 | 0.260 |
| Hiari | pack dog | 0.719 | 0.592 — 0.824 | **0.001** |
| Hakima | pet dog | 0.391 | 0.271 — 0.521 | 0.103 |
| Asali | pet dog | 0.375 | 0.257 — 0.505 | 0.060 |
| Freya | pet dog | 0.438 | 0.314 — 0.567 | 0.382 |
| Pepeo | pet dog | 0.500 | 0.372 — 0.628 | 1.000 |
| Zazu | pet dog | 0.328 | 0.216 — 0.457 | **0.008** |
| Coco | pet dog | 0.234 | 0.138 — 0.357 | **< 0.001** |
